# Cryo-EM structure of human LRRC15 reveals the basis of therapeutic antibody recognition

**DOI:** 10.64898/2026.06.16.732218

**Authors:** Xiaomin Wang, Mihin Perera, Maryam Sana, Chunxiao Wang, Ching-Seng Ang, Hariprasad Venugopal, Phillip Pymm, Wai-Hong Tham, Jeffrey J. Babon, Andrew Leis, Rhys Grinter, Shabih Shakeel

## Abstract

Leucine-rich repeat containing 15 (LRRC15) is a structural uncharacterised membrane protein, which is highly upregulated in cancer-associated fibroblasts and enriched in desmoplastic tumours including pancreatic, breast and head-and-neck cancers, where it is associated with therapy resistance. The LRRC15-targeting antibody-drug conjugate samrotamab vedotin has advanced into Phase I clinical evaluation, yet mechanistic understanding of how samrotamab engages its target has remained elusive in the absence of structural data. Here, using hydrogen-deuterium exchange mass spectrometry and cryo-electron microscopy, we define the samrotamab epitope on LRRC15 and determine the structure of the LRRC15-samrotamab complex to 2.6 Å resolution – the first high-resolution structure of this receptor. Samrotamab engages a membrane-proximal epitope within the C-terminal leucine-rich repeat region, leaving the canonical concave surface fully exposed for potential interactions. The lateral binding mode of samrotamab and the open arc geometry of LRRC15 provide a structural rationale for why antibody occupancy may leave signalling-competent surfaces intact. Guided by these insights, we computationally designed minibinders targeting the concave surface, identifying multiple binders with nanomolar affinity. Together, our work establishes the first structural framework for LRRC15, defines the molecular basis of therapeutic antibody recognition, and identifies the concave surface as a promising target for next-generation LRRC15-directed therapeutics.

## Main

Leucine-rich repeat containing 15 (LRRC15)^1^ is a type I transmembrane protein expressed in cancer-associated fibroblasts (CAFs) that plays a critical role in the tumour microenvironment^2^. Its restricted expression pattern and association with poor prognosis across multiple cancer types have positioned LRRC15 as an attractive therapeutic target for cancer immunotherapy^3^. In triple-negative breast cancer (TNBC), LRRC15 is highly upregulated and its depletion suppresses tumour cell proliferation, migration and invasion through inhibition of ITGB1/FAK/PI3K/AKT signalling pathways^4^. Beyond cancer, LRRC15 has been implicated in rheumatoid arthritis, where it is overexpressed in fibroblast-like synoviocytes and promotes inflammatory activation, synovial proliferation and bone destruction^5,6^. Silencing LRRC15 attenuates disease progression in collagen-induced arthritis models, highlighting a broader pathogenic role for LRRC15 in fibroblast-driven diseases. Together, these findings establish LRRC15 as a clinically relevant target across malignant and fibroblast-driven inflammatory diseases.

The development of LRRC15-targeted antibody-drug conjugates, particularly samrotamab vedotin (ABBV-085), represents a promising strategy to selectively deliver cytotoxic payloads to the tumour stroma while sparing normal tissues^2^. Samrotamab is a fully human IgG1 monoclonal antibody that specifically recognises human LRRC15 and has advanced into Phase I clinical trials, demonstrating preliminary antitumour activity in subsets of sarcoma patients^7^. Despite growing therapeutic interest, no high-resolution structure of LRRC15 has been determined, limiting mechanistic understanding of antibody engagement, receptor architecture and ligand recognition. Understanding the structural basis of antibody-antigen recognition is essential for rational therapeutic development, including affinity optimisation, and the design of alternative binding modalities, particularly for membrane-proximal targets, where receptor topology and cell-surface presentation can strongly influence therapeutic efficacy.

Leucine-rich repeat (LRR) proteins form a large and structurally diverse superfamily found across all kingdoms of life, subdivided into distinct subfamilies including Typical, RI-like, Bacterial, SDS22-like and plant-specific, based on repeat length, sequence consensus and cellular context^8–10^. Among these, extracellular Typical LRR proteins adopt an open horseshoe-shaped solenoid in which parallel β-strands lining the concave face serve as the primary platform for protein–protein and protein–ligand interactions^11–13^, in contrast to intracellular RI-like proteins such as the ribonuclease inhibitor, which adopt a substantially more compact, closed-arc architecture; other subfamilies such as Bacterial and SDS22-like display distinct repeat geometries that reflect their varied functional and organismal contexts^8,12^. As a type I transmembrane protein bearing a predicted extracellular LRR domain, LRRC15 is expected to adopt the open Typical fold, making structural characterisation of its ectodomain central to understanding its receptor architecture and potential for therapeutic targeting.

Recent advances in cryo-electron microscopy (cryo-EM) have enabled high-resolution structure determination of antibody-antigen complexes, and computational protein design methods have opened routes to alternative binding scaffolds. Tools such as BindCraft^14^ leverage deep learning models to design de novo synthetic binders, or minibinders, targeting specific receptor surfaces in compact, stable formats that offer potential advantages in tissue penetration, immunogenicity and manufacturing compared with conventional antibodies.

Here, we integrate hydrogen–deuterium exchange mass spectrometry (HDX-MS), cryo-EM and computational protein design to determine the first high-resolution structure of LRRC15, define the molecular basis of samrotamab recognition and engineer minibinders to target a distinct surface on LRRC15.

## Results

### Samrotamab recognises a C-terminal epitope on human LRRC15

To characterise samrotamab recognition of LRRC15, we recombinantly expressed the human LRRC15 ectodomain (residues 1-470) in insect cells and purified samrotamab from mammalian cells (**Supplementary Fig. 1**). Fab fragments were generated by IgdE protease digestion and incubated with LRRC15 at a 3:1 molar excess prior to purification by size-exclusion chromatography (SEC). SEC analysis showed a clear shift in elution volume consistent with stable complex formation, confirmed with SDS–PAGE showing co-elution of LRRC15 and samrotamab^Fab^ (**Fig. 1a** and **Supplementary Fig. 1d**).

**Figure 1.**
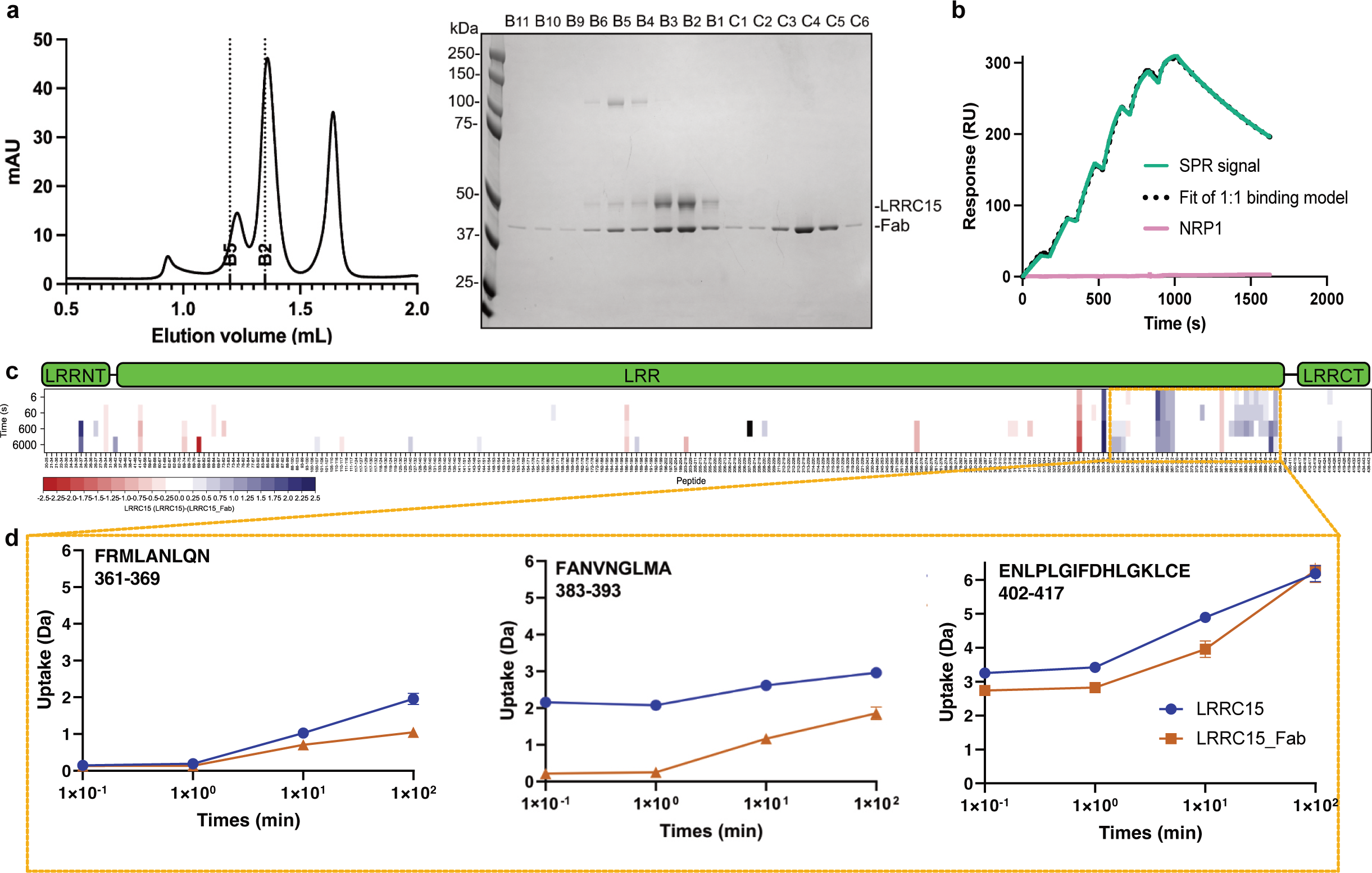
Mapping the samrotamab epitope on LRRC15. **a.** Purification of the LRRC15-samrotamab^Fab^ complex. Size-exclusion chromatography (left) of LRRC15 incubated with samrotamab^Fab^ at a 1:3 molar ratio run on Superdex200 Increase 3.2/300 column (Cytiva). Coomassie-stained SDS–PAGE analysis (right) of indicated SEC fractions of the LRRC15-samrotamab^Fab^ complex in non-reducing conditions. M, molecular weight marker. **b.** Surface plasmon resonance (SPR) analysis of LRRC15 binding to samrotamab. LRRC15 was injected over a Protein A sensor chip captured with samrotamab antibody using a two-fold dilution series with the highest concentration at 250 nM. Sensorgrams were fitted using a 1:1 binding model. NRP1 protein was used as a negative control at concentrations up to 100 μM (pink). K_D_, equilibrium dissociation constant is 5.42 ± 0.39 nM; k_a_, association rate constant is (1.37 ± 0.04) ×10□ M ^-1^s^-1^ and; k_d_, dissociation rate constant is (7.43 ± 0.32) × 10^□4^ s^-1^. Error bars represent range from independent replicate experiments (n = 2). **c.** Hydrogen–deuterium exchange mass spectrometry (HDX-MS) analysis of LRRC15 in the absence and presence of samrotamab^Fab^. Chicklet plot showing statistically significant differences in deuterium uptake upon complex formation. Regions protected upon binding are shown in blue, and exposed regions in red. Colouring indicates a difference in uptake of ≥ 5% with P ≤ 0.01 (Welch’s one-sided t-test; n = 3 independent experiments). The domain diagram of LRRC15 is shown above the chicklet plot. LRRNT is Leucine Rich Repeat N-terminus, LRR is Leucine Rich Repeat, and LRRCT is Leucine Rich Repeat C-terminus. **d.** Representative HDX-MS deuterium uptake plots for LRRC15 peptides in the absence (blue) or presence (red) of samrotamab^Fab^ for the regions likely to be protected on Fab binding. Error bars represent mean ± 2 standard deviations from technical replicates (n = 3).

We next quantified the interaction between LRRC15 and samrotamab using surface plasmon resonance (SPR). LRRC15 bound samrotamab with high affinity, yielding an equilibrium dissociation constant (K_D_) of ∼5.42 nM, with rapid association and slow dissociation kinetics (**Fig. 1b**). The unrelated receptor neuropilin-1 (NRP1) showed no detectable binding, confirming the specificity of the interaction. Together, these data establish that samrotamab engages LRRC15 with high affinity and forms a stable complex suitable for biophysical and structural studies.

To map the samrotamab epitope on LRRC15, we performed HDX-MS comparing apo LRRC15 with the LRRC15-samrotamab^Fab^ complex. HDX-MS achieved 95.5% sequence coverage with high peptide redundancy, enabling near-complete interrogation of the LRRC15 ectodomain (**Supplementary Fig. 2 and Supplementary Table 1**). Differential deuterium uptake analysis identified strongly protected regions towards the C-terminal end of the LRR domain upon Fab binding (**Fig. 1c**). Representative uptake plots demonstrated substantial protection within residues 361–369, 383–393 and 402–417 in the presence of samrotamab^Fab^, consistent with direct antibody engagement of this region (**Fig. 1d**).

To validate these HDX-defined epitope, we generated a series of LRRC15 truncation mutants and assessed samrotamab binding by SPR (**Supplementary Fig. 3**). Deletion of residues distal to the LRR C-terminal cap (Δ426–470) had minimal effect on binding affinity relative to wild-type LRRC15, indicating that this tail region does not contribute substantially to antibody recognition. In contrast, progressive truncations removing epitope 3 (Δ402–470), epitopes 2–3 (Δ385–470), or all three epitopes (Δ360–470) abolished detectable binding. Together, these data demonstrate that samrotamab recognises a conformational epitope formed by clustered LRR elements within the C-terminal region of LRRC15.

### Cryo-EM structure of LRRC15-samrotamab^Fab^

To establish the structural basis of LRRC15 recognition and determine the first high-resolution structure of this receptor, we determined the cryo-EM structure of the LRRC15-samrotamab^Fab^ complex at 2.63 Å overall resolution (**Fig. 2a**, **Supplementary Fig. 4 and Supplementary Table 2**). The cryo-EM map exhibited well-resolved side-chain density across most of the complex, enabling de novo model building of the Fab and confident modelling of LRRC15 residues 24–426 (**Supplementary Fig. 5**). Residues 1–23 and 427–470 were not resolved, consistent with conformational flexibility in the N-terminal region and membrane-proximal tail.

**Figure 2.**
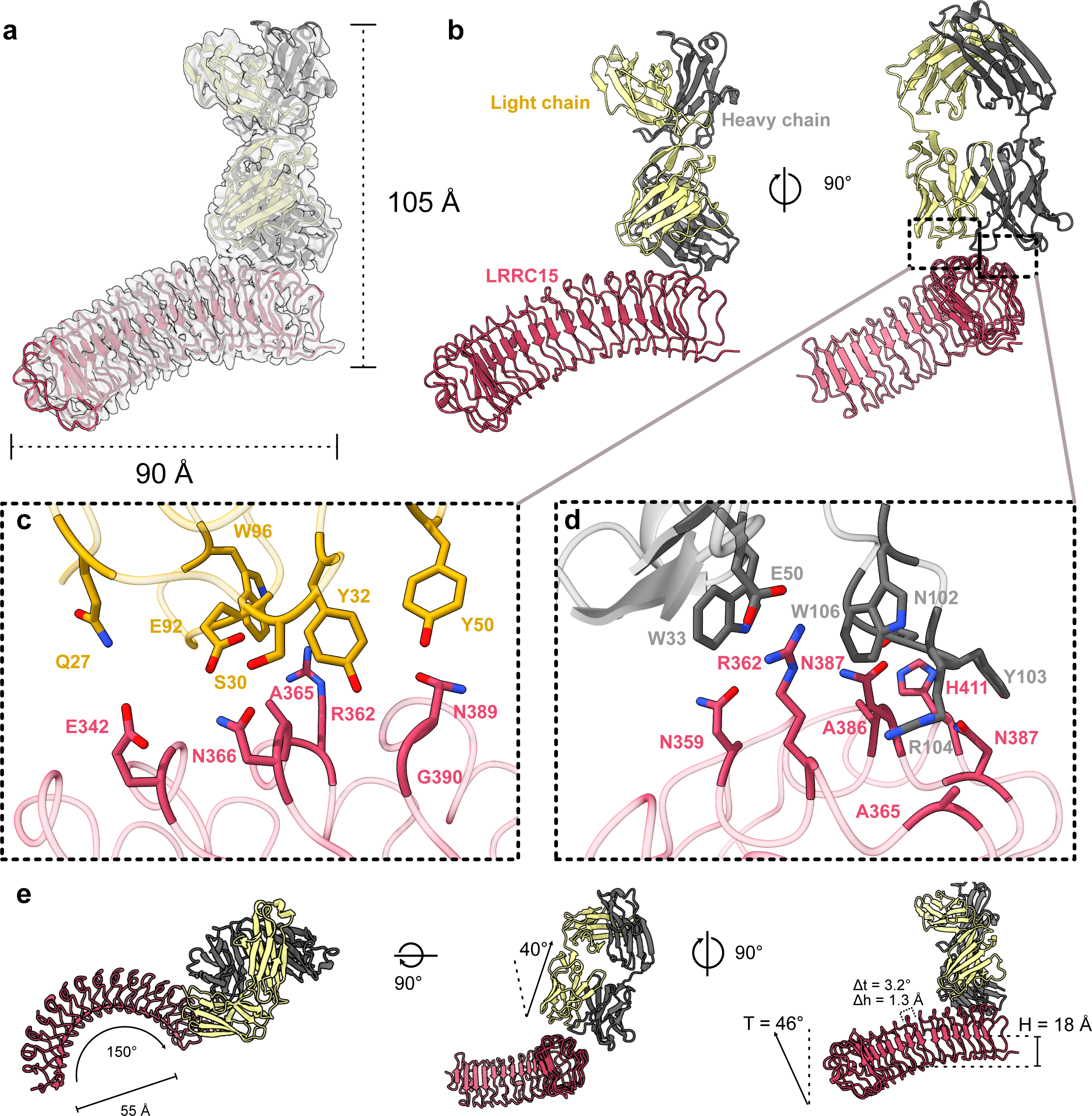
Cryo-EM structure of the LRRC15-samrotamab^Fab^ complex. **a-b**. Cryo-EM reconstruction and atomic model of the LRRC15-samrotamab^Fab^ complex shown in multiple orientations. LRRC15 is shown in pink, the samrotamab^Fab^ heavy chain in dark grey and the light chain in pale yellow. **c**. Close-up view of the interaction interface between the light chain of samrotamab^Fab^ and LRRC15. Residues participating in intermolecular interactions are shown as sticks and labelled. **d**. Close-up view of the interaction interface between the heavy chain of samrotamab^Fab^ and LRRC15. Most intermolecular contacts are mediated through the heavy chain of complementarity-determining regions (CDRs), including residues W33, Y103, R104 and W106. LRRC15 residues identified by HDX-MS are shown in pink. **e**. Geometric analysis of the LRRC15 LRR domain. The LRRC15 ectodomain forms a pronounced right-handed superhelix with an overall arc of ∼150° and an internal diameter of ∼55 Å. Samrotamab^Fab^ engages LRRC15 at an angle of ∼40° relative to the longitudinal axis of the LRR domain. Side and top views illustrate the curvature and orientation of the complex.

Samrotamab^Fab^ binds the C-terminal LRR region of LRRC15, in agreement with the HDX-MS data (**Fig. 2b-d**). The antibody engages the outer edge of the LRR scaffold at the interface between the canonical concave β-sheet surface and the convex outer surface – a binding mode that contrasts with the concave-surface engagement typically observed for extracellular LRR receptors^11,15^. Samrotamab instead recognises a composite epitope centred on loop regions connecting the β-strands and short helical elements of the C-terminal LRR repeats, approaching at approximately 40° relative to the longitudinal axis of the ectodomain (**Fig. 2e middle panel**). Interface analysis showed that the majority of the buried surface area is contributed by the heavy chain (472 Å²) relative to the light chain (357 Å²), with most intermolecular contacts formed by heavy-chain complementarity-determining regions (CDRs) residues N102, Y103 and R104 (**Supplementary Tables 3 and 4**). Comparison of the cryo-EM interface with the HDX-MS data demonstrated strong agreement: apart from two peripheral residues (N359 and E342), all LRRC15 residues directly contacting the Fab localised within the three protected HDX epitopes regions.

The LRRC15 ectodomain adopts the characteristic crescent-shaped architecture of LRR proteins, comprising an elongated curved solenoid of 16 canonical repeats (residues 38–397) flanked by N- and C-terminal capping regions. Geometric analysis using an in-house developed package called banana-roll (β-strand and arc-based normal-plane analysis of near-planar angles in repeating local LRR geometry package), placed LRRC15 firmly within the open-arc branch of the extracellular Typical LRR subfamily^12^, with a fitted arc radius of ∼29.8 Å and an internal diameter of ∼55 Å – substantially more open than intracellular RI-like LRR proteins (∼17–20 Å) but consistent with extracellular receptors such as decorin, slit and Nogo receptor family members (**Fig. 2e, Supplementary Fig. 6 and Supplementary Table 5**). A small mean axial rise per repeat (∼1.3 Å) combined with a progressive increase in β-strand tilt across successive repeats generates a gently twisted continuous β-sheet surface reminiscent of the Möbius-strip-like topology described for other extracellular LRR proteins, particularly pronounced in LRRC15 owing to its unusually long repeat array (**Supplementary Fig. 6c**)^12^.

Critically, samrotamab targets a C-terminal lateral epitope while leaving the concave β-sheet surface fully exposed – a structural arrangement with direct implications for understanding both the mechanism of antibody engagement and the surfaces available to binding partners.

### Rational design of concave surface-targeting LRRC15 minibinders

Since ligand recognition in many leucine-rich repeat proteins occurs predominantly through the concave β-sheet surface^8–10,13^, we hypothesised that this region may represent a functionally important interaction platform on LRRC15 and therefore an attractive target for alternative therapeutic binders.

We used the BindCraft computational protein design platform to generate minibinders targeting the concave surface spanning the N-terminal to middle region of the LRRC15 ectodomain (**Fig. 3**). A total of 48 candidate minibinders were experimentally evaluated (**Supplementary Fig. 7**), from which five binders – H6, H3, F3, H2 and A5, displayed reproducible nanomolar binding affinity for LRRC15 as determined by SPR (**Fig. 3a-e, Supplementary Table 6**). Sensorgrams for H6, F3 and H2 were well described by a 1:1 binding model, consistent with specific monovalent interactions. In contrast, H3 and A5 exhibited binding behaviour that was not adequately captured by a 1:1 kinetic model and were therefore analysed using a steady-state affinity model. Among the five binders, H6 exhibited the highest affinity for LRRC15 (K_D_ = 179 ± 2 nM), followed by H3 (231 ± 17.7 nM) and F3 (283 ± 14.1 nM), whereas H2 and A5 displayed affinities of approximately 400–500 nM (**Fig. 3a-e**).

**Figure 3.**
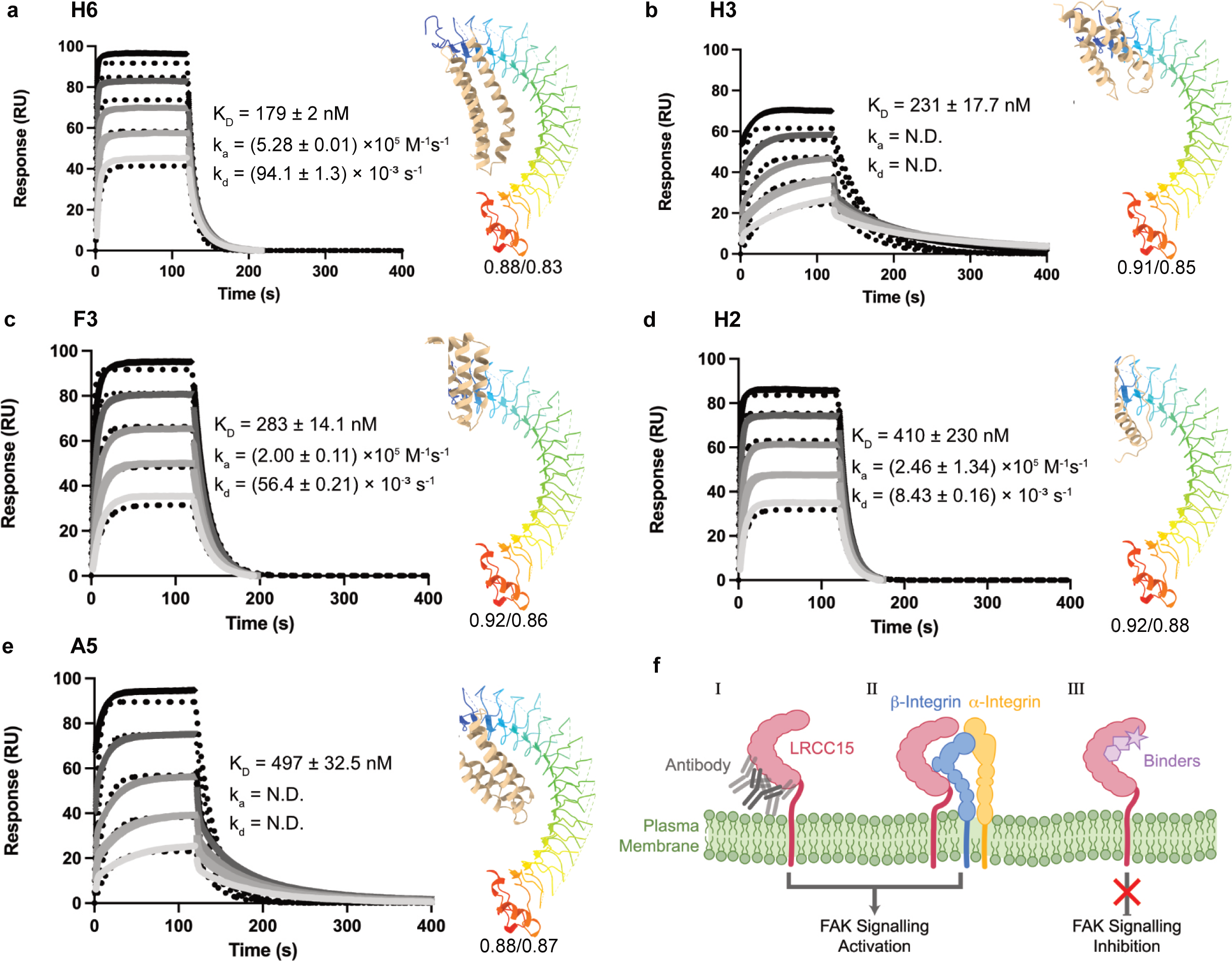
High affinity minibinders targeting the concave surface of LRRC15. Five high-affinity minibinders generated using the BindCraft computational design platform and validated by SPR: **a**, H6; **b**, H3; **c**, F3; **d**, H2; and **e**, A5. Minibinders were designed to target the N-terminal to the middle region of LRRC15 and bind exclusively to the canonical concave surface of the LRR domain. SPR sensorgrams were obtained by injecting a two-fold dilution series of minibinders over sensor chips captured with LRRC15, with the highest concentration of 2 μM. Solid lines represent experimental binding data, and dotted lines represent global fits to a 1:1 binding model. Equilibrium dissociation constants (K_D_), association rate constants (k_a_) and dissociation rate constants (k_d_) are indicated for each minibinder. Error values represent range from replicate experiments (n = 2). Structural models generated by BindCraft and predicted using AlphaFold2 are shown adjacent to the corresponding sensorgrams. LRRC15 is displayed as a cartoon coloured by rainbow spectrum from N-terminus (blue) to C-terminus (red), whereas minibinders are shown in single solid colours. The pTM/ipTM score for each predicted structure is shown below the model. All selected minibinders engage in the concave surface of LRRC15, spatially distinct from the samrotamab epitope. **f.** Proposed models for LRRC15-mediated signalling, antibody engagement and blocking. Schematic illustrating three possible modes of LRRC15 function at the plasma membrane. (I) Samrotamab binds a lateral epitope on LRRC15 while leaving the canonical concave LRR surface exposed. (II) The accessible concave surface may facilitate interaction with endogenous signalling partners, including integrins, potentially supporting downstream FAK signalling despite antibody occupancy. (III) Therapeutic binders targeting the concave surface may sterically block endogenous ligand interactions and inhibit downstream signalling pathways.

Computational structural models generated by BindCraft^14^ based on AlphaFold2^16^ predictions showed that all five minibinders engage the concave surface of LRRC15, occupying regions spatially distinct from the samrotamab epitope (**Fig. 3a-e, Supplementary Data 1**). The predicted complexes were supported by high-confidence AlphaFold2 metrics (pTM = 0.88–0.92; ipTM = 0.83–0.88), consistent with reliable prediction of both the overall complex architecture and the relative placement of the minibinders on the LRRC15 surface. While experimental confirmation of the precise binding location will require further structural characterisation, the models consistently position the minibinders across a broad region of the inner β-sheet face that is canonically used for ligand recognition in extracellular LRR proteins. These findings suggest that the concave surface of LRRC15 is targetable by computationally designed binders and provide a structural foundation for developing next-generation LRRC15-directed therapeutics that may directly interfere with endogenous signalling interactions (**Fig. 3f**).

## Discussion

LRRC15 has emerged as an important therapeutic target in both cancer and inflammatory disease owing to its highly restricted expression pattern and pathogenic roles in fibroblast-associated signalling pathways. Although the LRRC15-targeting antibody–drug conjugate samrotamab vedotin demonstrated an acceptable safety profile and preliminary antitumour activity in Phase I evaluation, broader clinical benefit across advanced solid tumours was limited^7^. Here, by integrating HDX-MS and cryo-EM, we establish the first structural framework for LRRC15 and reveal molecular features that provide a direct structural rationale for these observations.

Our structure reveals that samrotamab engages the lateral edge of the LRR solenoid, at the interface between the concave β-sheet surface and the outer convex face, within the C-terminal portion of the structured LRR domain. Our cryo-EM model resolves LRRC15 residues 24–426; however, the full extracellular domain extends to residue 538, meaning that 112 residues connecting the structured LRR C-terminal cap to the transmembrane domain remain uncharacterised. The precise orientation and flexibility of this stalk region, and therefore the absolute distance between the samrotamab epitope and the cell membrane, cannot be determined from the current structure. Nonetheless, a further consideration is that affinity measurements using soluble recombinant ectodomains lacking this stalk and membrane context may not fully recapitulate antibody engagement on intact cells, where glycosylation, receptor clustering and the physical constraints of the cell surface could collectively alter binding dynamics and accessibility. Such factors may be particularly limiting within the dense extracellular matrix environments of desmoplastic tumours. Together, these factors suggest caution in extrapolating solution-phase affinity measurements to the cell surface and highlight the importance of evaluating therapeutic ADC engagement in membrane-presented formats.

A notable feature of the structure is that samrotamab binds along the lateral edge of the LRR solenoid while leaving the entire canonical concave β-sheet surface exposed. This binding mode differs markedly from ligand engagement in many extracellular LRR receptors, where the concave surface serves as the primary interaction platform^8–10^. The open arc geometry of LRRC15 and the preservation of an accessible concave surface suggest that endogenous signalling interactions may remain intact despite antibody occupancy, an observation potentially relevant to understanding the limits of samrotamab’s clinical activity.

These structural observations are particularly significant given emerging evidence that LRRC15 contributes directly to oncogenic signalling rather than acting solely as a stromal marker. In TNBC, LRRC15 depletion suppresses proliferation, migration and invasion through inhibition of ITGB1/FAK/PI3K/AKT signalling^4^, while in rheumatoid arthritis, LRRC15 promotes inflammatory activation of fibroblast-like synoviocytes and synovial destruction^5,6^. The large accessible concave surface and open arc geometry of LRRC15 are consistent with canonical ligand-recognition modes of extracellular LRR proteins^12,17^ and support the hypothesis that LRRC15 engages endogenous signalling partners, potentially including integrins, through this surface. An antibody that occupies only the lateral edge of the molecule would therefore leave these interactions intact, offering a structural explanation for why samrotamab’s clinical benefit was confined to cytotoxic payload delivery rather than signalling modulation.

These insights motivated our targeting of the concave surface with computationally designed minibinders. Unlike conventional antibodies, compact synthetic scaffolds may provide improved penetration into dense tumour microenvironments and access to sterically restricted epitopes. The identification of five nanomolar-affinity minibinders predicted to engage the concave surface of LRRC15 at a site spatially distinct from the samrotamab epitope establishes a proof-of-principle that this canonical LRR interaction surface is selectively targetable by engineered binders. Whether concave-surface engagement modulates LRRC15-mediated signalling remains to be established, and determining the functional consequences of minibinder binding including effects on integrin interactions and downstream FAK signalling represents an important priority for future work. Nevertheless, these findings provide a structural and computational foundation for next-generation therapeutic strategies that may simultaneously improve cell-surface accessibility and directly interfere with endogenous LRRC15 signalling.

Beyond its therapeutic implications, our work establishes the first experimentally determined high-resolution structure of LRRC15 and reveals an unusually open solenoidal architecture with a gently twisted concave surface that most closely approximates the Möbius-strip-like topology described for the extracellular Typical LRR family. More broadly, our findings highlight how epitope topology, rather than affinity alone, may critically determine the efficacy of stromal-targeting antibody therapeutics. Overall, we establish the structural basis for samrotamab recognition of LRRC15, identify a C-terminal lateral epitope whose cell-surface accessibility warrants further evaluation in membrane-presented contexts, and demonstrate the concave surface as a promising target for next-generation LRRC15-directed therapeutic development in cancer and inflammatory disease.

## Methods

### Protein expression and purification

Ectodomain of human LRRC15 (UniProt id: Q8TF66, 1-470) cDNAs were synthesized by GenScript Biotech. For protein expression, LRRC15 was sub-cloned into a pACEBac1 vector with a C-terminal 3 C protease-cleavage site followed by Twin StrepII-tag. DNA construct was transformed into EMBacY cells to generate a recombinant bacmid. Recombinant bacmid DNA was transfected into Sf21 cells using FuGENE transfection reagent (Promega) and virus was passaged once (P2) in the same cell line before large-scale infection. Sf21 cells were a kind gift from Yibin Wu, Lawrence lab, WEHI (negative for mycoplasma). For large-scale expression, Sf21 cells at a density of 1 × 10^6^ cells per mL were infected with 1% (v/v) P2 virus and incubated at 27 °C for 60 h, 220 rpm in a shaker incubator.

The culture was harvested 2.5 days post-infection and supernatant was collected after centrifugation. LRRC15 protein was eluted from streptactin XT beads (IBA) in buffer containing 25 mM Tris pH 8.5, 300 mM NaCl and 50 mM biotin. The elution fraction containing LRRC15 was concentrated using a 10 kDa molecular weight cut-off concentrator (Merck Millipore) and further purified by size exclusion chromatography (SEC) using the Superdex 75 10/300 GL column (Cytiva).

### Expression, purification of samrotamab and generation of its Fab

The gene sequences encoding the variable heavy (VH) and variable light (VL) chains of the therapeutic monoclonal antibody samrotamab (patent id: WO 2021/067673 A1) were codon-optimised for expression in mammalian cells. The genes were synthesized *in vitro* and subsequently cloned into mammalian vector pcDNA3.4 (GenScript Biotech).

For transient expression, Expi293F™ cells (Thermo Fisher Scientific) were maintained in Expi293™ Expression Medium at 37 °C, 8% CO₂, and 125 rpm in a shaking incubator. Cells were seeded at a density of 2.5 x 10⁶ viable cells/mL and were transfected with plasmids encoding the heavy and light chains using polyethylenimine (PEI, 1 mg/mL, pH 7.0) at a 1:3 DNA-to-PEI mass ratio.

The cell culture supernatant was harvested 4 days post-transfection by centrifugation at 4,000 × *g* for 30 minutes to remove cells and debris. The clarified cell culture supernatant was filtered through a 0.22 µm polyethersulfone (PES) membrane to remove any residual particulates. The filtrate was immediately loaded onto a 1 mL HiTrap MabSelect PrismA™ protein A column (Cytiva) pre-equilibrated with 3 column volumes (CV) of filtered binding Buffer (1X Dulbecco’s phosphate-buffered saline, DPBS, pH 7.5). The column was subsequently washed with 10 CV of filtered Binding Buffer to remove unbound contaminants. Bound antibodies were eluted using a buffer comprising 0.1 M citric acid, pH 3.0. The eluted fractions were collected into tubes containing 150-200 µL 1 M Tris-HCl (pH 9.0) per mL elution volume for immediate neutralisation to pH 7.5 to preserve antibody integrity. The antibody was concentrated using an Amicon® Ultra-15 centrifugal filter (50 kDa MWCO, Millipore Sigma) and loaded onto a Superdex 200 Increase 10/300GL SEC column (Cytiva) pre-equilibrated with 25 mM HEPES pH 7.5, 150 mM NaCl. The peak corresponding to the monomeric antibody was collected.

Fab fragments were generated by enzymatic cleavage of whole IgG using cysteine protease IgdE. Briefly, IgdE was added to the IgG at an enzyme-to-substrate (E:S) ratio of 1:20 (w/w). The reaction mixture was incubated at 37 °C for overnight. The digest mixture was centrifuged at 16,000 xg to remove precipitates and immediately loaded onto a 1 mL HiTrap™ KappaSelect column (Cytiva) pre-equilibrated with 3 column volumes (CV) of filtered binding Buffer (1X Dulbecco’s phosphate-buffered saline, DPBS, pH 7.5). The column was washed with 10 CV of filtered Binding Buffer to remove unbound contaminants.

Bound Fab was eluted using 0.1 M glycine, pH 1.5. The eluted fractions were immediately neutralised by adding one-tenth volume of 1 M Tris-HCl (pH 9.0) to preserve Fab integrity. The Fab was concentrated using an Amicon® Ultra-15 centrifugal filter (10 kDa MWCO, Millipore Sigma) and loaded onto a Superdex 75 Increase 10/300GL SEC column (Cytiva) pre-equilibrated with 25 mM HEPES pH 7.5, 150 mM NaCl. The peak corresponding to the monomeric Fab was collected.

### HDX-MS data collection and data processing

Human LRRC15 and samrotamab^Fab^ were mixed (1:2 molar ratio) at 4□ O/N and subjected to size exclusion chromatography using Superdex 200 Increase 10/300GL Colum (Cytiva). The fractions corresponding to the peak of complex were collected and concentrated to final concentration of 25 µM. HDX labelling of protein was performed at 20 °C for periods of 0, 6, 60, 600, and 6000 sec using a PAL Dual Head HDX Automation manager (Trajan/LEAP) controlled by the ChronosHDX software. 3 µL of the purified protein (∼25 µM) sample was transferred to 55 µL of non-deuterated (50 mM potassium phosphate buffer pH 8 in water) or deuterated (50 mM potassium phosphate buffer pD 7.6 in D_2_O) buffer and incubated for the respective time. Quenching was performed by adding 50 µL of the deuterated protein to 50 µL of quench buffer (50 mM potassium phosphate buffer, pH 2.3, 4 M guanidine hydrochloride and 0.1% n-dodecylphosphocholine, 200 mM TCEP) at 1 °C. For online pepsin digestion, 80 µL of the quenched sample was passed over an immobilised 2.1 × 30 mm Enzymate BEH pepsin column (Waters) at 15°C, equilibrated in 0.1% formic acid in water at 100 µL/min. To further reduce peptide carryovers, n-octyl-β-d-glucopyranoside at 1% w/w was added the pepsin column wash solution (1.5M guanidine hydrochloride, 4% acetonitrile, 0.8% formic acid, 150 mM TCEP). Proteolysed peptides were captured in cooling chamber maintained at 1°C and desalted by a C18 trap column (VanGuard BEH; 1.7 μm; 2.1 × 5 mm; (Waters)) and eluted with acetonitrile and 0.1% formic acid gradient (5% to 40% in 8 min, 40% to 95% in 0.5 min, 95% 1.5 min) at a flow rate of 80 μL/min and separated on an ACQUITY UPLC BEH C18 analytical column (1.7 μm, 1 × 50 mm, (Waters) delivered by ACQUITY UPLC I-Class Binary Solvent Manager (Waters).

For mass spectrometry, an ion mobility equipped SYNAPT G2-Si mass spectrometer (Waters) was used. Instrument settings were: 3.0 kV capillary and 40 V sampling cone with source and desolvation temperature of 100 and 40 °C, respectively. The desolvation and cone gas flows were at 800 L/hr and 100 L/hr, respectively. High energy ramp trap collision energy was from 20 to 40 V. All mass spectra were acquired using a 0.4 sec scan time with continuous lock mass (Leu-Enk, 556.2771m/z) for mass accuracy correction. Data were acquired in HDMSE (ion mobility) mode, and peptides from non-deuterated samples were identified using Protein Lynx Global Server (PLGS) v3.0 (Waters). Primary digest was set to non-specific with methionine oxidation as the variable modification. The FDR was set at 1%. To ensure high peptide selection stringency, we applied additional filter constraints of 0.3 fragments per residue, minimum consecutive product of 1, minimum intensity of 1000, maximum MH+ error of 10 ppm, retention time RSD of 10% and file threshold of 2 out of 3 HDMSE files. The deuterium uptake values were calculated for each peptide using DynamX 3.0 (Waters). The average back-exchange in our system (∼30%) was measured using phosphorylase B, but we did not adjust for back exchange as our analysis compares relative deuteration between protein states, and therefore all results are reported as relative deuterium exchange levels expressed in mass units (Da). Deuterium exchange experiments were performed in triplicate for each of the timepoints. Peptides with a statistically significance difference in HDX were determined using the HD-eXplosion software with a p-value ≤ 0.01 in a Welch’s t-test (n = 3)^18^.

### Cryo-EM sample preparation and data collection

LRRC15 was incubated with samrotamab^Fab^ at a 1:3 molar ratio and run on Superdex200 Increase 3.2/300 column (Cytiva). The fraction containing the LRRC15-samrotamab^Fab^ complex was vitrified by applying 4 μL purified complex (0.13 mg/ml), to UltrAuFoil R1.2/1.3 grids (Quantifoil). The grids were glow discharged at 100% power for 3 mins using a 200 W glow discharge instrument (Henniker Plasma). The grids were prepared using a Vitrobot Mark IV (Thermo Fisher) at 4 °C and 100% humidity with arbitrary blotting force of 0 N for 5 s. The data were collected with EPU software on a Krios G4 transmission electron microscope operating at 300 keV using a 4K x 4K Falcon 4i detector (Thermo Fisher).

### Image Processing

LRRC15-samrotamab^Fab^ complex was processed in CryoSPARC v4.7.0^19^ (**Supplementary Fig. 4**). 21,866 movies were imported, motion-corrected and CTF estimated. Micrographs with CTF fit greater than 5 Å, and full frame motion greater than 200 pixels were discarded, leaving 16,274 micrographs. Particles were picked with PartiNet^20^ on micrographs denoised with the integrated PartiNet denoiser. 2,948,409 particle coordinates were imported into CryoSPARC, and particles were extracted with a box size of 284 pixels and Fourier downsampled to 192 pixels. These particles were initialised with “*ab initio* Reconstruction” with 4 classes. The classes were then heterogeneously refined. The two classes representing the complex were selected for “Non-Uniform Refinement” with a user-generated mask. Particles were then extracted at 284 pixels, with updated 2D and 3D aligned shifts. Iterative “Non-Uniform Refinement” was performed, optimising per particle scale, particle defocus and per-beam shift Global CTF aberrations (Tilt, Trefoil, Tetrafoil, Spherical Aberration and Anisotropic Magnification). Particles were then extracted at 320 pixels, with updated 2D and 3D aligned shifts, and further iterative “Non-Uniform Refinement” with per-particle scale and CTF optimisation was performed. This resulted in an initial map of 2.57 Å with 1,762,077 particles.

Inspection of this map revealed anisotropy in the Fab and tip of the LRRC15 density, due to undersampling of views along the short axes of the complex. Templates were generated and selected from this initial volume corresponding to the missing views, and template picking was performed with a low pass filter of 6 Å on selected templates. These particles were extracted with a box size of 284 pixels and Fourier downsampled to 80 pixels. Two rounds of 2D classification with 150 classes were used to filter junk particles and overrepresented views. Iterative “Non-Uniform Refinement” was performed on these particles against the initial volume, optimising per-particle scale and defocus. Particles were then extracted at 320 pixels, with updated 2D and 3D aligned shifts. Another round of “Non-Uniform Refinement” was performed with the full resolution particles. These particles were combined with the stack from the initial map, and “Rebalance Orientations” was performed with rebalance percentile = 80% and exclusion criteria = 3Dalignments/alpha. This resulted in a final combined stack of 1,957,225 particles. Finally, “Local Refinement” was performed, with gold-standard half splits redone and the particle pose/shift gaussian prior to alignment. This resulted in the final consensus map of 2.63 Å with stronger orientation metrics.

### Model building

An initial model for the Fab fragment was built using ModelAngelo v1.01^21^ against a locally refined cryo-EM map, with a mask around the Fab. Missing residues and loops were subsequently completed using phenix.fit_loops^22^. For the LRRC15 component, an AlphaFold2^14^ predicted model was docked into the cryo-EM density using phenix.predict_and_build, followed by manual trimming of poorly fitting loop regions.

The Fab and LRRC15 models were combined into a single complex model in UCSF ChimeraX v1.10^23^ and rigid body fitted into the full complex map. The combined model was subjected to real-space refinement using phenix.real_space_refine. Iterative manual model building and validation was performed in ISOLDE v1.10^24^ within UCSF ChimeraX, with Ramachandran outliers, rotamer corrections, and steric clashes resolved iteratively. Secondary structure restraints were applied throughout ISOLDE refinement to preserve helical and strand geometry. Model geometry was assessed after each refinement round using MolProbity^24^. Cryo-EM data collection, reconstruction, refinement, and model building statistics are provided in **Supplementary Table 2**.

### Structural geometry analysis of the LRRC15 LRR domain

A three-dimensional geometric analysis of the LRRC15 leucine-rich repeat (LRR) domain was performed. Sixteen canonical repeats were defined along the inner β-strand of the LRR stack, excluding cap repeats. For each repeat, the reference Cα was at the strand midpoint (position 4); β-strand orientation was defined by Cα atoms at positions 3 and 5. Position-4 Cα residues spanned residues 59–418 (mean spacing 24 residues per repeat).

Sixteen anchor coordinates were used to fit two global models: a three-dimensional circle (arc radius and total arc) and a parametric superhelix (least-squares fit with multi-start L-BFGS-B optimisation). Between consecutive anchors, three local step metrics were calculated: chord distance, the straight-line Cα–Cα distance from one repeat to the next; axial rise, the component of that step along the fitted superhelix axis (advance along the long axis of the fitted superhelix); and curvature, the angle by which the path bends between successive steps along the arc. β-Sheet tilt was quantified for each repeat as the angle between the strand direction vector (Cα at position 5 minus Cα at position 3) and the plane of the best-fit circle through all position-4 anchors—i.e. how much each inner β-strand deviates from lying orthogonal to the plane of the fitted circle. Per-repeat values were summarised as mean ± standard deviation across the sixteen repeats and global descriptors included total arc, internal diameter *D* (= 2*R* sin(φ[/2) from the circle fit), mean axial rise, and mean β-sheet tilt are provided **(Supplementary Table 5**). Analysis was performed with banana-roll v0.1 (developed in-house) available at https://github.com/MihinP/banana-roll written in Python 3.14 with Gemmi, NumPy, and SciPy. Geometry plots were prepared in R 4.5.1.

### In silico minibinders generation by Bindcraft

Synthetic protein binders were designed using BindCraft^14^. To enhance computational efficiency, a trimmed AlphaFold2^16^ model of LRRC15 encompassing only the target-relevant region was used as the structural input. The standard BindCraft workflow was applied without modification of scoring custom filters. Designs were guided by hotspot-directed targeting of a concave surface on LRRC15 to bias sampling toward geometrically and chemically complementary binding modes. All binders were constrained to 60–100 amino acids (8–14 kDa).

For each target site, the top 24 candidates were selected based on automated BindCraft scoring (∼100 total designs). Final sets were refined by manual inspection, excluding models lacking a well-formed hydrophobic core or exhibiting non-physical or unstable folds.

### Expression and purification of minibinders

Minibinder gene sequences were synthesised by Twist Bioscience and inserted in a pET29b expression vector between NdeI and XhoI binding sites with a C-terminal 6xHis tag. A total of 48 top-ranked minibinder constructs were expressed in *E. coli* C41 (DE3) cells using the Overnight Express Autoinduction System (Overnight Express^TM^ Instant TB Medium, Merck 71491-4). Cultures were prepared in 50mL Terrific Broth (TB) medium supplemented with kanamycin and distributed into 125 mL conical flasks. Each flask was inoculated with 500 μL of starter culture and incubated at 30 °C for 16–24 h with shaking.

Cells were harvested by centrifugation, and pellets were weighed and stored at −20 °C until further use. For lysis, cell pellets were resuspended in B-PER Bacterial Protein Extraction Reagent (Thermo Scientific, 78248) using 5 mL reagent per gram of wet cell mass, supplemented with lysozyme (chicken egg white powder, Sigma-Aldrich 62971-10G-F) and DNase (bovine pancreas grade II, Roche 10104159001) according to the manufacturer’s instructions. Lysis was performed at room temperature for 15 min with agitation. Insoluble material was removed by centrifugation at 4,000 xg for 15 min at 4 °C, and clarified lysates were collected.

Purification was performed by immobilised metal affinity chromatography (IMAC) under batch-binding conditions. Clarified lysates were incubated with 100 μL nickel-agarose resin per sample in a 24-well plate at 4 °C for 2 h with shaking. The resin was transferred to a 24-well filter plate, and unbound material was removed by vacuum filtration. The resin was washed three times with 5 mL of nickel binding buffer (50 mM Tris, 200 mM NaCl, 20 mM Imidazole, pH 7.9). Bound proteins were eluted twice with 500 μL nickel elution buffer (50 mM Tris, 200 mM NaCl, 500 mM Imidazole, pH 7.9), each for 10 min at room temperature.

### Surface Plasmon Resonance (SPR)

All SPR binding studies were performed using a Biacore 8K Instrument (Cytiva). The binding studies were performed at 18 °C in SPR running buffer. The sensorgrams were double referenced, and steady-state binding data were fitted using a 1:1 binding model using Biacore 8K Evaluation Software (Cytiva).

LRRC15 and samrotamab: Samrotamab was diluted to 1 µg/mL in SPR running buffer (10 mM HEPES pH 8, 150 mM NaCl, 3 mM EDTA and 0.005% (v/v) Tween-20) to a final immobilization level of 200-300 response units (RU) on the Protein A sensor chip (Cytiva). A blank activation/deactivation was used for the reference surface. LRRC15 was diluted to 250 nM in SPR running buffer and prepared as a 6-point concentration series (2-fold serial dilution, 8.7-250 nM). Samples were injected in a single cycle run (flow rate 30 µL/min, contact time of 120 s, dissociation 600 s). Protein Neuropilin 1 ectodomain (NRP1) was used as a negative control.

LRRC15 deletion constructs and samrotamab: Samrotamab was diluted to 0.5 µg/mL in SPR running buffer to a final immobilization level of 200-220 RU on the Protein A sensor chip (Cytiva). A blank activation/deactivation was used for the reference surface. Deletion constructs of LRRC15 were diluted to 31.25 nM in SPR running buffer and prepared as a 6-point concentration series (2-fold serial dilution, 0.98-31.25 nM). Samples were injected in a single cycle run (flow rate 30 µL/min, contact time of 120 s, dissociation 600 s).

LRRC15 and minibinders: LRRC15 was diluted to 2 µg/mL in SPR running buffer to a final immobilization level of about 100 response units (RU) on the Strep-tactin® XT-modified sensor chip (Xantec). A blank activation/deactivation was used for the reference surface. The 48 mini-binders were diluted to 2000 nM in SPR running buffer and prepared as a 5-point concentration series (2-fold serial dilution, 125-2000 nM). Samples were injected in a multi-cycle run (flow rate 30 µL/min, contact time of 120 s, dissociation 600 s).

## Data availability

The mass spectrometry data is available from ProteomeXchange Consortium via the PRIDE partner repository with identifier PXD078980. The Cryo-EM map has been deposited in the EM Data Bank with the following accession code: EMD- 80945. Atomic coordinates have been deposited in the Protein Data Bank with the accession code 26XD. The raw micrographs are deposited in EMPIAR (EMPIAR-13589).

## Code availability

The banana-roll package (v0.1.0; https://github.com/MihinP/banana-roll) implements LRR repeat geometry analysis (circle and superhelix fitting, per-repeat chord, axial rise, curvature, and β-sheet tilt) and is freely available and distributed under the MIT License.

## Acknowledgements

We acknowledge the use of transmission electron microscopes at Ian Holmes Imaging Centre, Bio21 Institute, University of Melbourne, and the Monash University Ramaciotti Centre for Cryo-Electron Microscopy. We thank the WEHI Cryo-EM Platform, the WEHI Research Computing Platform and Milton high-performance computing facility and the Monash University MASSIVE high-performance computing facility for providing facilities and support. We would like to thank the Melbourne Mass Spectrometry and Proteomics Facility of The Bio21 Molecular Science and Biotechnology Institute at The University of Melbourne for the support of mass spectrometry analysis. We thank Ethan Goddard-Borger for the kind gift of IgdE plasmid and Chloe Gerak for help with Figure 3f. XW is supported by ARC CCeMMP/WEHI PhD scholarship. MP is supported by the Research Training Program (RTP) scholarship from Faculty of Engineering and Technology, University of Melbourne and Graeme Clark Institute for Medical Engineering Top-Up scholarship. W-HT is supported by National Health and Medical Research Council of Australia (NHMRC) GNT2016908, and PP is supported by NHMRC GNT2021267. W-HT and PP acknowledge the Victorian State Government Operational Infrastructure Support and Australian Government NHMRC IRIISS. R.G. was funded by an NHMRC EL1 Investigator Grant (APP1197376) and the Grimwade Fellowship in Biochemistry (University of Melbourne). SS is supported by funds from WEHI, the estate of Akos and Marjorie Talon, The University of Melbourne Attraction and Retention Funds and the NHMRC Investigator grant (GNT2016827).

## Author contributions

XM performed protein purification, sample preparation for HDX-MS, SPR, and cryo-EM. XM and CA performed HDX-MS and its data analysis. XM and JJB performed SPR. XM, HV and AL screened and collected cryo-EM data. XM and MP performed cryo-EM image analysis. MP performed model building, refinement, and geometry parameter measurements. RG designed minibinders. CW expressed and purified minibinders. MS performed cellular assays. XM and SS wrote the manuscript with input from all the co-authors. PP and WHT contributed towards experimental design and data interpretation. SS conceived and supervised the project.

## Competing interests

The authors declare no competing interests

## Notes

### Competing Interest Statement

The authors have declared no competing interest.

## References

1 Satoh, K., Hata, M. & Yokota, H. A novel member of the leucine-rich repeat superfamily induced in rat astrocytes by beta-amyloid. Biochemical and biophysical research communications 290, 756–762 (2002). 10.1006/bbrc.2001.6272

2 Purcell, J. W. et al. LRRC15 Is a Novel Mesenchymal Protein and Stromal Target for Antibody-Drug Conjugates. Cancer research 78, 4059–4072 (2018). 10.1158/0008-5472.CAN-18-0327

3 Ray, U. et al. Exploiting LRRC15 as a Novel Therapeutic Target in Cancer. Cancer research 82, 1675–1681 (2022). 10.1158/0008-5472.CAN-21-3734

4 Wu, X. et al. Leucine rich repeat containing 15 promotes triple-negative breast cancer proliferation and invasion via the ITGB1/FAK/PI3K signalling pathway. Scientific reports 15, 14535 (2025). 10.1038/s41598-025-98661-1

5 Xin, M., Xia, G., Guan, X., Xi, G. & Fu, M. Leucine-Rich Repeat Containing 15 Promotes the Inflammatory Response in Rheumatoid Arthritis by Regulating NF-kappaB Pathway. Immun Inflamm Dis 13, e70220 (2025). 10.1002/iid3.70220

6 Ding, H. et al. RUNX1 Ameliorates Rheumatoid Arthritis Progression through Epigenetic Inhibition of LRRC15. Mol Cells 46, 231–244 (2023). 10.14348/molcells.2023.2136

7 Demetri, G. D. et al. First-in-Human Phase I Study of ABBV-085, an Antibody-Drug Conjugate Targeting LRRC15, in Sarcomas and Other Advanced Solid Tumors. Clin Cancer Res 27, 3556–3566 (2021). 10.1158/1078-0432.CCR-20-4513

8 Kajava, A. V. Structural diversity of leucine-rich repeat proteins. Journal of molecular biology 277, 519–527 (1998). 10.1006/jmbi.1998.1643

9 Kobe, B. & Deisenhofer, J. Proteins with leucine-rich repeats. Current opinion in structural biology 5, 409–416 (1995). 10.1016/0959-440x(95)80105-7

10 Buchanan, S. G. & Gay, N. J. Structural and functional diversity in the leucine-rich repeat family of proteins. Prog Biophys Mol Biol 65, 1–44 (1996). 10.1016/s0079-6107(96)00003-x

11 Kobe, B. & Kajava, A. V. The leucine-rich repeat as a protein recognition motif. Current opinion in structural biology 11, 725–732 (2001). 10.1016/s0959-440x(01)00266-4

12 Enkhbayar, P., Kamiya, M., Osaki, M., Matsumoto, T. & Matsushima, N. Structural principles of leucine-rich repeat (LRR) proteins. Proteins 54, 394–403 (2004). 10.1002/prot.10605

13 Song, W. et al. Structural basis for specific recognition of single-stranded RNA by Toll-like receptor 13. Nature structural & molecular biology 22, 782–787 (2015). 10.1038/nsmb.3080

14 Pacesa, M. et al. One-shot design of functional protein binders with BindCraft. Nature 646, 483–492 (2025). 10.1038/s41586-025-09429-6

15 Brodsky, I. & Medzhitov, R. Two modes of ligand recognition by TLRs. Cell 130, 979–981 (2007). 10.1016/j.cell.2007.09.009

16 Jumper, J. et al. Highly accurate protein structure prediction with AlphaFold. Nature 596, 583–589 (2021). 10.1038/s41586-021-03819-2

17 Matsushima, N., Enkhbayar, P., Kamiya, M., Osaki, M. & Kretsinger, R. H. Leucine-Rich Repeats (LRRs): Structure, Function, Evolution and Interaction with Ligands. Drug Design Reviews - Online (Discontinued) 2, 305–322 (2005). 10.2174/1567269054087613

18 Zhang, N., Yu, X., Zhang, X. & D’Arcy, S. HD-eXplosion: visualization of hydrogen-deuterium exchange data as chiclet and volcano plots with statistical filtering. Bioinformatics 37, 1926–1927 (2021). 10.1093/bioinformatics/btaa892

19 Punjani, A., Rubinstein, J. L., Fleet, D. J. & Brubaker, M. A. cryoSPARC: algorithms for rapid unsupervised cryo-EM structure determination. Nature methods 14, 290–296 (2017). 10.1038/nmeth.4169

20 Perera, M. et al. PartiNet is a dynamic adaptive neural network for high-performance particle picking in cryo-electron microscopy. bioRxiv, 2026.2001.2023.700950 (2026). 10.64898/2026.01.23.700950

21 Jamali, K. et al. Automated model building and protein identification in cryo-EM maps. Nature 628, 450–457 (2024). 10.1038/s41586-024-07215-4

22 Afonine, P. V. et al. Real-space refinement in PHENIX for cryo-EM and crystallography. Acta Crystallogr D Struct Biol 74, 531–544 (2018). 10.1107/S2059798318006551

23 Meng, E. C. et al. UCSF ChimeraX: Tools for structure building and analysis. Protein science : a publication of the Protein Society 32, e4792 (2023). 10.1002/pro.4792

24 Croll, T. I. ISOLDE: a physically realistic environment for model building into low-resolution electron-density maps. Acta Crystallogr D Struct Biol 74, 519–530 (2018). 10.1107/S2059798318002425

