## Supplementary Information for "Cryo-EM structure of human LRRC15 reveals the basis of therapeutic antibody recognition"

**Supplementary information for**
### Cryo-EM structure of human LRRC15 reveals the basis of therapeutic antibody recognition


Xiaomin Wang[1,2], Mihin Perera[1,3], Maryam Sana[1], Chunxiao Wang[4,5], Ching-Seng Ang[4], Hariprasad Venugopal[5], Phillip Pymm[1,6], Wai-Hong Tham[1,6], Jeffrey J. Babon[1,6], Andrew Leis[1,6], Rhys Grinter[4,5], Shabih Shakeel[1,2,4*]

[1]The Walter and Eliza Hall Institute of Medical Research, 1G Royal Parade, Parkville, Melbourne, VIC 3052, Australia

[2]ARC Centre for Cryo-electron Microscopy of Membrane Proteins, Bio21 Molecular Science and Biotechnology Institute, The University of Melbourne, Parkville, Melbourne, VIC 3052, Australia

[3]Department of Chemical Engineering, The University of Melbourne, Parkville, Melbourne, VIC 3052, Australia

[4]Department of Biochemistry and Pharmacology, The University of Melbourne, Parkville, Melbourne, VIC 3052, Australia

[5]Department of Microbiology, Biomedicine Discovery Institute, Monash University, Clayton, Melbourne, VIC 3168, Australia

[6]Department of Medical Biology, The University of Melbourne, Parkville, VIC 3052, Melbourne,

Australia



Keywords: cryo-EM, LRRC15, protein design, cancer, antibodies

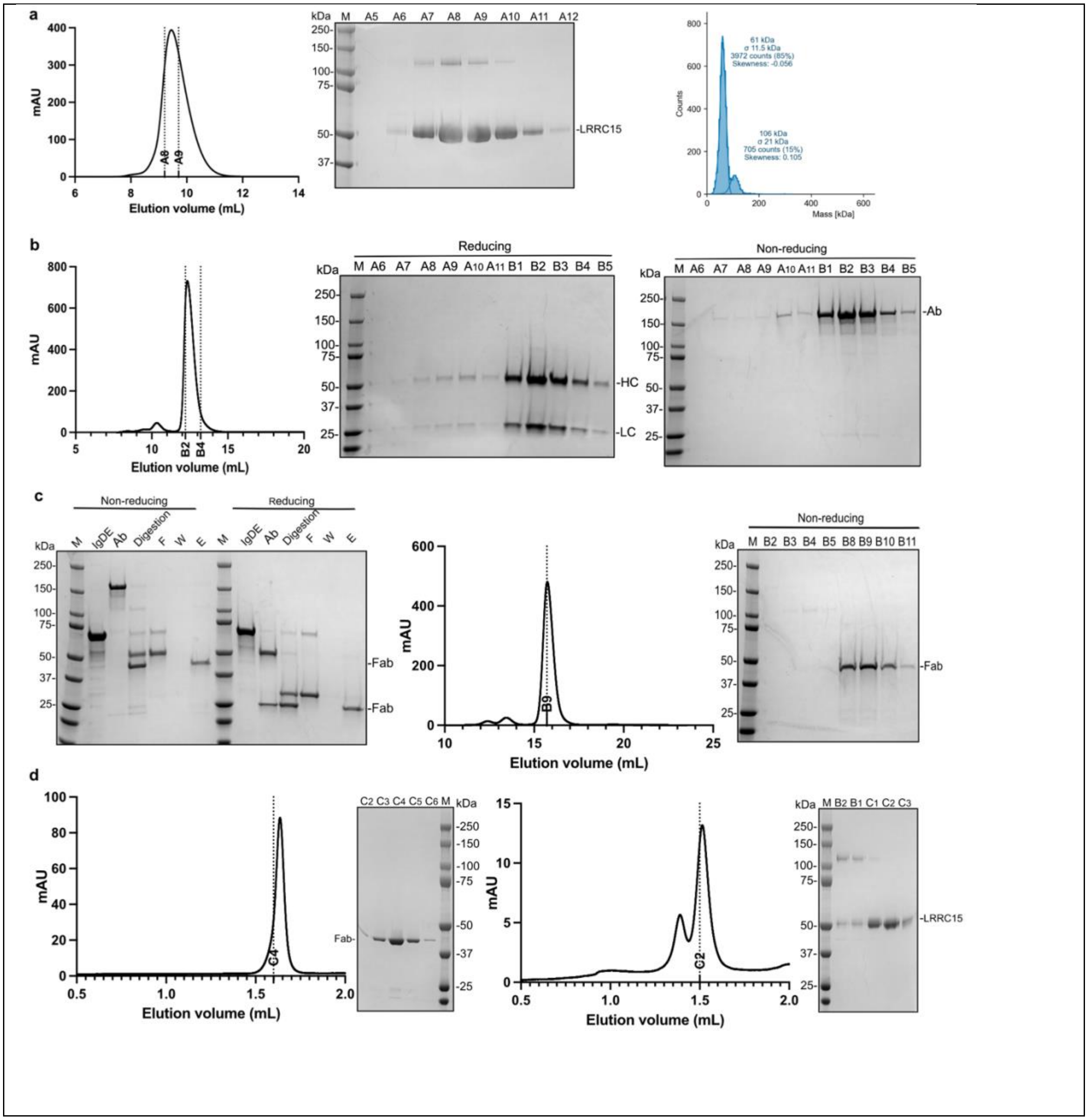


***Supplementary Figure 1. Purification of LRRC15, samrotamab and samrotamab$^{Fab}$.***

**a**. Purification of the human LRRC15 ectodomain. Size-exclusion chromatography (SEC) profile of recombinant LRRC15 ectodomain (residues 1–470) expressed in insect cells and purified by affinity and SEC chromatography (left). Peak fractions corresponding to the major SEC peak were analysed by non-reducing SDS–PAGE and visualized by Coomassie staining (middle). Mass photometry showing that LRRC15 ectodomain purifies predominantly as a monomer (right). **b**. Purification of recombinant samrotamab IgG1. SEC profile of full-length samrotamab antibody expressed in Expi293F cells and purified using Protein A affinity chromatography followed by SEC (left). Reducing SDS–PAGE analysis (middle) shows separation of heavy chain (HC) and light chain (LC), whereas non-reducing SDS–PAGE analysis (right) confirms the integrity of assembled antibody (Ab). **c**. Generation and purification of samrotamab$^{Fab}$. SDS-PAGE analysis of IgdE-mediated digestion of samrotamab under reducing and non-reducing conditions (left). SEC profile of purified Fab following affinity purification and size-exclusion chromatography (middle). Non-reducing SDS-PAGE analysis of SEC peak fractions confirms purification of monomeric Fab fragments (right). M, molecular weight marker; F, flowthrough; W, wash; E, elution. **d**. SEC profile and SDS-PAGE analysis of samrotamab$^{Fab}$ (left) and LRRC15 (right) from Superdex200 Increase 3.2/300 column (Cytiva).

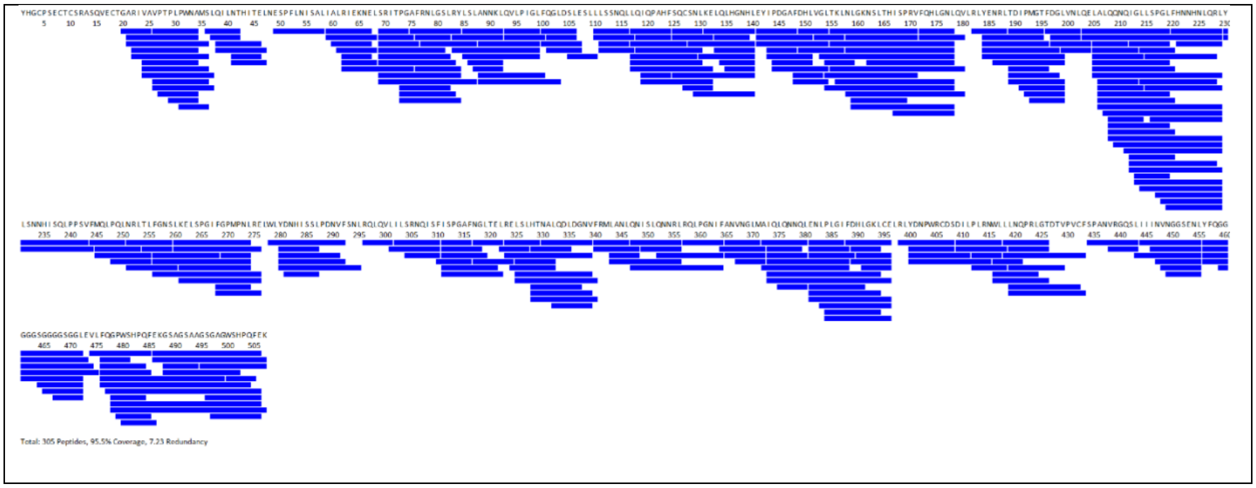


***Supplementary Figure 2. HDX-MS sequence coverage map of LRRC15.***

A total of 305 peptides (average redundancy for covered amino acids of 7.23) were identified, representing 95.5% sequence coverage of LRRC15.

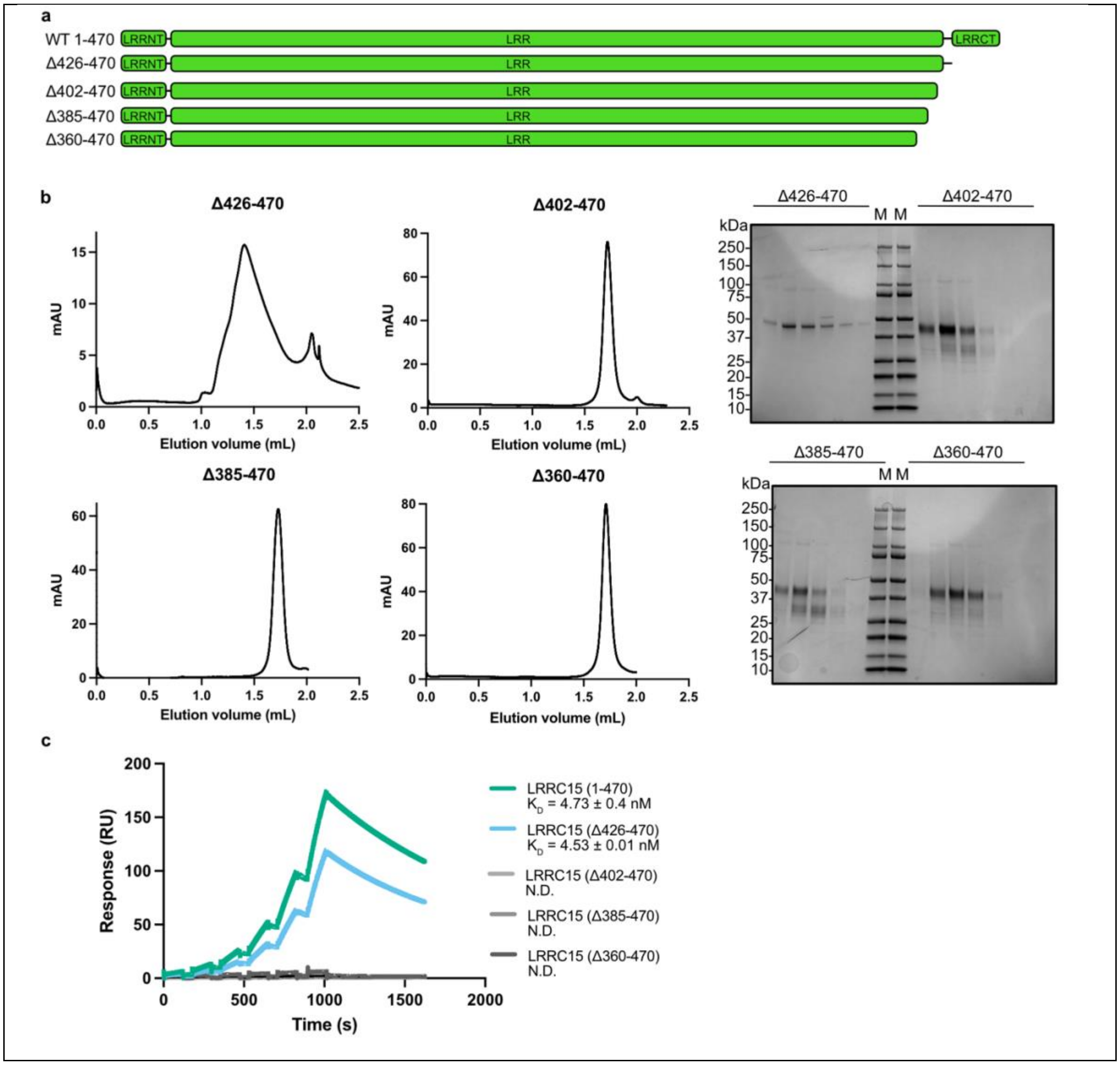


***Supplementary Figure 3. Deletion of epitope(s) but not C terminal disrupts the LRRC15 and samrotamab interaction.***

**a**. Deletion constructs design for LRRC15 Δ426-470 (C-terminal deletion), LRRC15 Δ360-470 (LRR14-Cterminus end), LRRC15 Δ385-470 (LRR14-Cterminus end) and LRRC15 Δ402-470 (LRR15-C terminus end). **b**. Purification of the human LRRC15 deletion constructs. SEC profile for each construct is shown on left. Peak fractions corresponding to the major SEC peak were analysed by non-reducing SDS–PAGE and visualized by Coomassie staining for each construct (right). M, molecular weight marker. **c**. SPR binding curve of WT LRRC15 ECD, Δ426-470, Δ360-470, Δ385-470 and Δ402-470. The experiment was repeated three independent times (n=3). A two-fold dilution series of antigen LRRC15 was injected over the chip surface; the highest concentration was 31.25 nM. $K_D$ means equilibrium constant. N.D. is not determined as there was no binding detected.

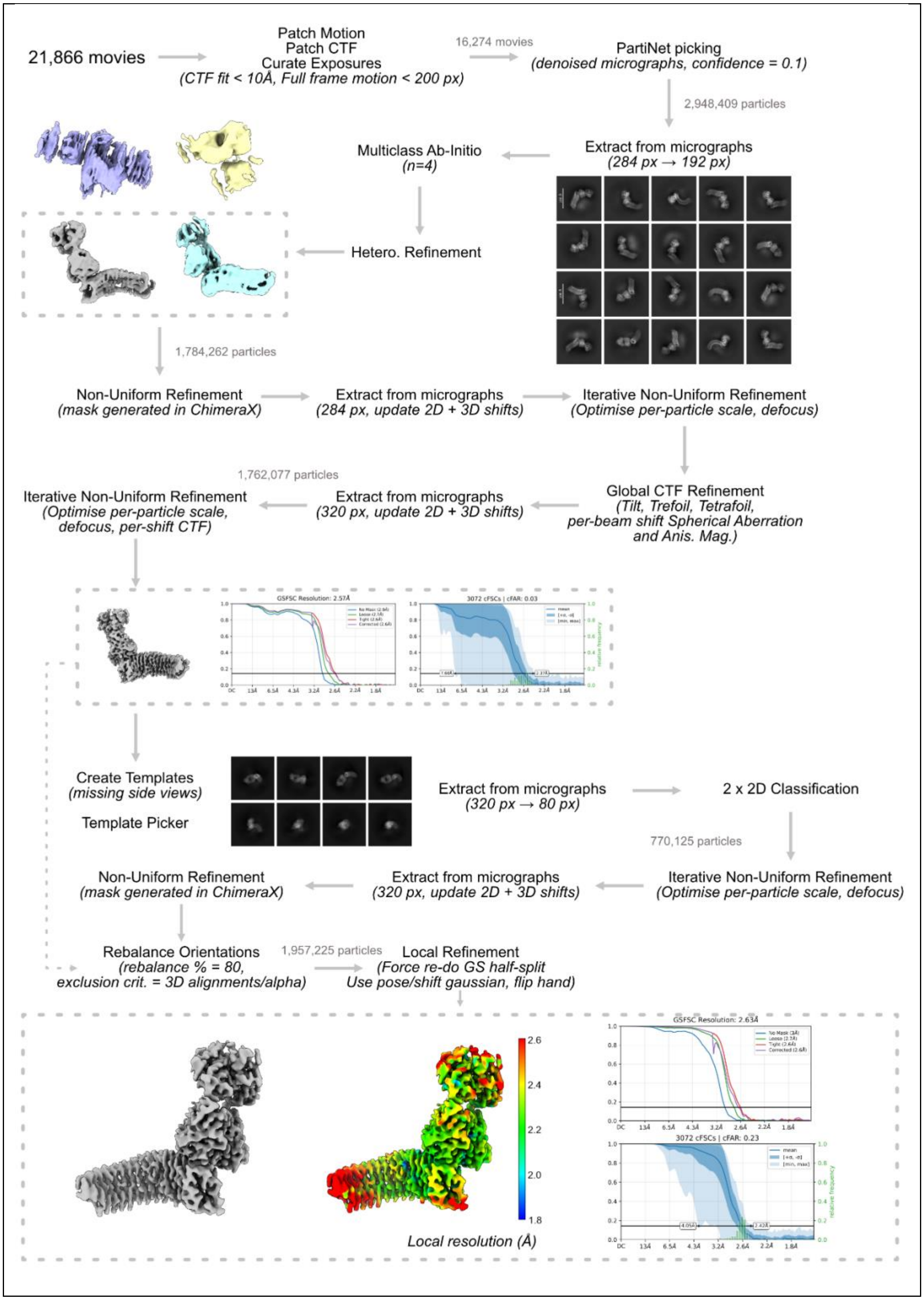


***Supplementary Figure 4. Cryo-EM image processing workflow for the LRRC15-samrotamab$^{Fab}$ complex.***

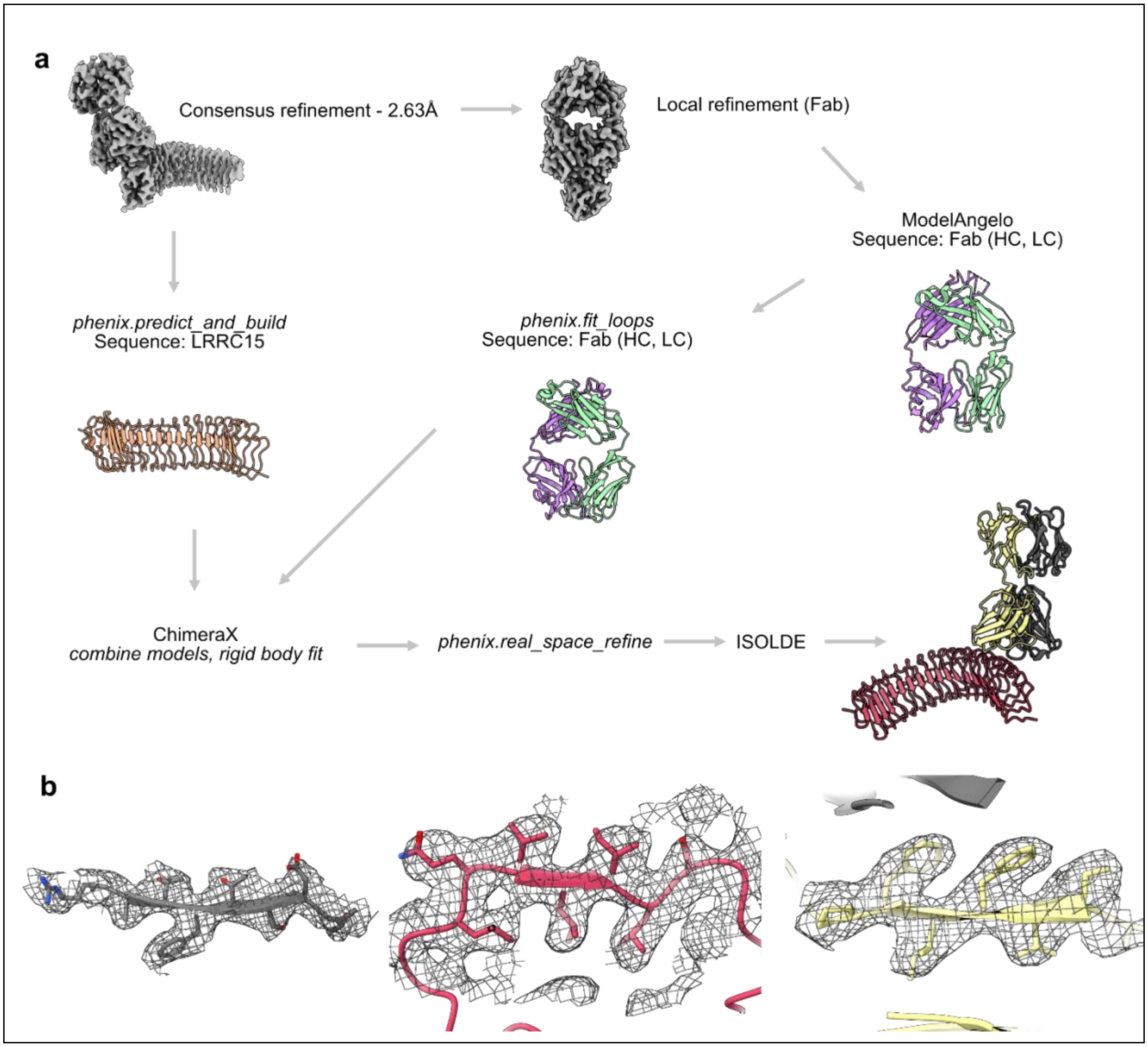


***Supplementary Figure 5. Model building and refinement for LRRC15-samrotamab$^{Fab}$ complex.***

**a**. Workflow showing modelling into LRRC15-samrotamab$^{Fab}$. **b**. Representative regions from LRRC15 and heavy and light chains of Fab showing map-model fit quality.

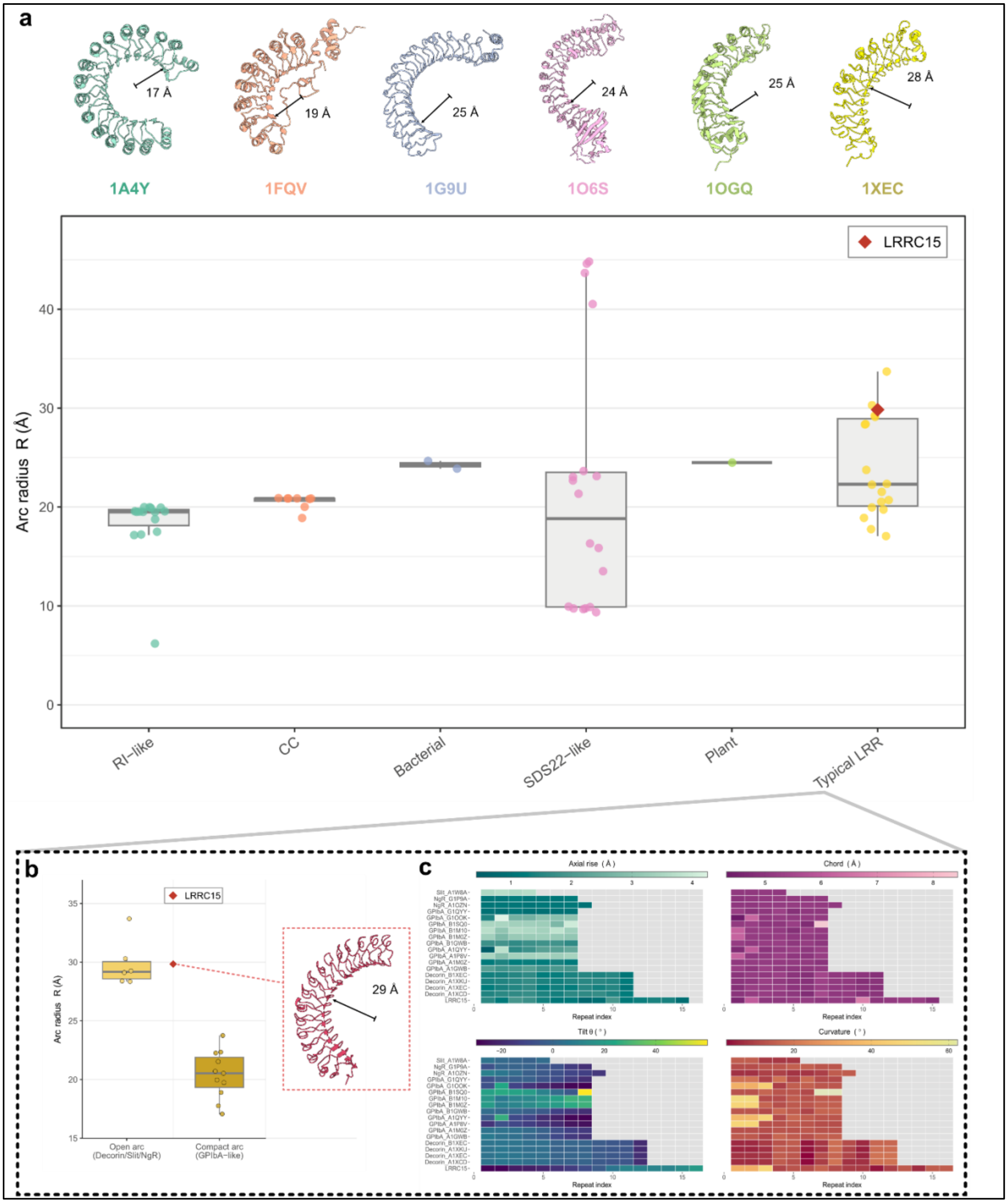


***Supplementary Figure 6. Geometric comparison of LRRC15 and other LRR family proteins.***

**a**. Global arc radius (R) across LRR families. Banana-roll circle fits to inner beta-strand anchors. Above each family column, a representative structure (RI-like: angiogenin inhibitor 1A4Y chain A; CC: Skp2 1FQV chain C; bacterial: YopM 1G9U chain A; SDS22-like: InlA 1O6S chain A; plant: PGIP 1OGQ chain A; Typical LRR: decorin 1XEC chain A is shown with its fitted arc radius R (Å) annotated. Box plots and points show computed R for all chains in that family; the red diamond is LRRC15 ($R \approx 29.8$ Å). **b**. Computed R for Typical LRR homologs grouped as open arc (Decorin, Slit, NgR; $R \geq 27$ Å) or compact arc (GPIbA-like; $R < 27$ Å), with LRRC15 (red diamond) shown separately. LRRC15 structure with fitted arc radius in red **c**. Heatmaps (protein × repeat index) for axial rise, chord, tilt θ, and curvature from banana-roll analysis for Typical LRR homologs and LRRC15 (caps excluded). Each row is one structure; colour scales are independent per metric.

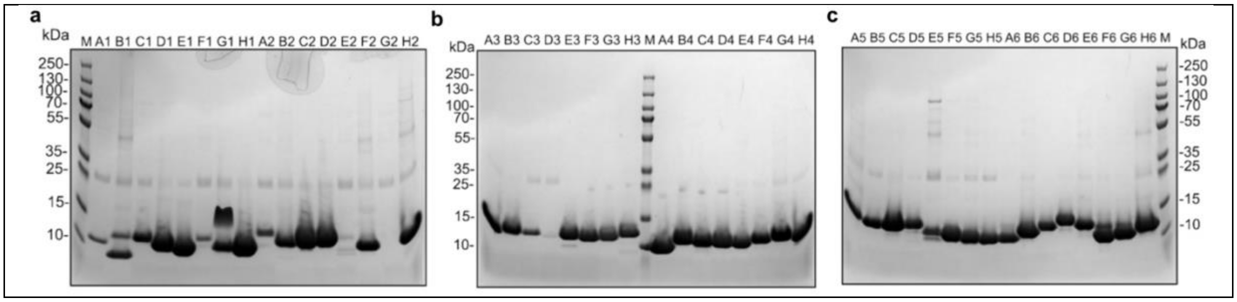


***Supplementary Figure 7. Expression and purification of minibinders.***

Coomassie stained non-reducing SDS-PAGE of minibinders after His-tag purification. (**a**) minibinders A1-H2 eluates, (**b**) A3-H4 eluates, (**c**) A5-H6 eluates. M: molecular weight protein markers.

*Supplementary Table 1. HDX-MS summary table*

| **Experimental Conditions** | **LRRC15, LRRC15-samrotamab$^{Fab}$** |
|---|---|
| HDX reaction details | 20 mM HEPES 4-(2-hydroxyethyl)-1-piperazineethanesulfonic acid, 150 mM NaCl, $pH_{read}$ = 7.6 |
| HDX time course (s) | 0, 6, 60, 600,6000 sec at 20 °C |
| HDX control samples | unlabeled LRRC15 or LRRC15 complex with Fab |
| Replicates (technical) | 3 |
| **LRRC15 vs LRRC15-samrotamab$^{Fab}$** | |
| # of peptides | 305 |
| Sequence coverage | 95.5% |
| Average peptide length / Redundancy | 7.23 |
| Calculated ΔHDX significance threshold | 0.31 Da / 2.69% |
| Mean Back-exchange (% of possible deuterium uptake) | 30% (measured using phosphorylase B) |

*Supplementary Table 2. Cryo-EM data collection, refinement and validation statistics*

| **Data Collection** | **LRRC15-samrotamab$^{Fab}$ (EMD- 80945 and EMPIAR-13589)** |
|---|---|
| Micrographs | 21,866 |
| Particles (Final map) | 1,957,225 |
| Pixel size (Å) | 0.808 |
| Defocus range (µm) | 0.4 – 1.4 |
| Voltage (kV) | 300 |
| Electron dose (e/Å$^2$) | 40 |
| Symmetry imposed | C1 |
| Initial particle images (no.) | 2,948,409 |
| Final particle images (no.) | 1,957,225 |
| Resolution (0.143 FSC) (Å) | 2.63 |
| Final Refinement | Local Refinement with GS-half splits in cryosparc |
| Map sharpening *B* | -89.6 |
| Model used | de novo using ModelAngelo and AlphaFold |
| Unmodelled regions | 1-23 and 427-470 |
| $CC_{map\ model}$ | 0.81 |
| **Model quality** | |
| Bond length (Å) / Bond angles (°) | 0.012/2.039 |

| **Ramachandran** | |
|---|---|
| Favoured (%) | 93.82 |
| Outliers (%) | 0.12 |
| Rotamer outliers (%) | 0.0 |
| C-Beta deviations (%) | 0.52 |
| Clashscore | 0.54 |
| MolProbity Score | 1.10 |

***Supplementary Table 3. Intermolecular residue–residue interactions between LRRC15 and Heavy chain of samrotamab$^{Fab}$***

| LRRC15 residue | Samrotamab$^{Fab}$ Heavy chain residue |
|---|---|
| N359* | W33 |
| R362 | W33, E50, R104, W106 |
| A365 | R104 |
| A386 | N102, Y103 |
| N387 | N102, Y103, R104, W106 |
| D410 | Y103 |
| H411 | N102, Y103 |

**Residues not identified as part of epitopes by HDX-MS*

***Supplementary Table 4. Intermolecular residue–residue interactions between LRRC15 and Light chain of samrotamab$^{Fab}$***

| LRRC15 residue | Samrotamab$^{Fab}$ Light chain residue |
|---|---|
| E342* | Q27 |
| R362 | W96 |
| A365 | Y32, E92 |
| N366 | S30, Y32, E92 |
| N389 | Y32, Y50 |
| G390 | Y32 |

**Residues not identified as part of epitopes by HDX–MS*

***Supplementary Table 5. LRRC15 geometric analysis statistics***

| **Parameter** | **Value** | **Method** |
|---|---|---|
| Number of canonical repeats | 16 | Manual selection of inner β-strand repeats |
| Repeat length | 24 residues | Mean spacing of position-4 Cα anchors |
| Residue span (canonical) | 59–418 | Position-4 Cα anchors (positions 3 and 5 for strand vectors) |
| Arc radius | 29.8 Å | 3D circle fit (Enkhbayar et al., 2003) |
| Total arc length | 147° | 3D circle fit |
| Superhelix radius | 28.1 Å | Global superhelix fit (multi-start L-BFGS-B) |
| Total arc (superhelix fit) | 153° | Global superhelix fit |
| Radius of curvature | 29.6 Å | Global superhelix fit |
| Curvature per repeat | 14.8° ± 11.3° | Local geometry (angle between successive anchor steps) |
| Chord distance per repeat | 5.23 ± 0.51 Å | Local geometry (3D Cα–Cα step, position 4) |
| Axial rise per repeat | 1.30 ± 0.34 Å | Local geometry (step projected onto superhelix axis) |
| Axial rise per repeat (global fit) | 1.17 Å | Global superhelix fit |
| Total axial rise | ~18 Å | Global superhelix fit |
| Superhelical twist per repeat | 3.1° ± 4.4° (right-handed) | β-strand vector rotation (Enkhbayar et al., 2003) |
| Total superhelical twist | ~46° | Sum of per-repeat β-strand twist |
| Circle fit RMSD | 1.37 Å | 3D circle fit (position-4 anchors) |
| Superhelix fit RMSD | 1.27 Å | Global superhelix fit (position-4 anchors) |

***Supplementary Table 6. Details of high affinity minibinders***

| **Minibinder** | **Sequence** |
|---|---|
| H6 | ASMIAEQYDKLLEVEEIKEEFEIEKELYEYRLEEARDNPEEFAKVKEEMQKIVEKMEELVKEVAAQRDASPALVKRMEEYVKEVEEMVKELVDKYTG |
| H3 | SYVEKVEEMKKKYESSKEHKPEELDEFLDFVVEMAEKFNLLDDPIFQEIMRHLILAKALYPEEGEDPEWNKELIEEHIKKAFELALEFAKKVDELEEKE |
| F3 | IEEELDKFYELLKKVAKENDLDIEKIREAMTPVVESFEDPRVRRAAQFMLDSVIGLLEFFARDGIKDENVVDWVYDWAKSHPEEFLRYAEERYPDL |
| H2 | SKAEELKAKGEEYWNKYYYLLELREEGKDPRSEEEAEAEINAAFSEAMTYINAYERLNPNGEKFVFLGSSEKPK |
| A5 | SEVEEWFEKLKEVIEKAISKFDPLFQSMIREEIFELFEERYREIKERVAAGDGVPLSVLMQQFIDELIEEFEFYSLLSEENKKELPKIIEELESL |
